# A continental system for forecasting bird migration

**DOI:** 10.1101/293092

**Authors:** Benjamin M. Van Doren, Kyle G. Horton

## Abstract

Billions of animals cross the globe each year during seasonal migrations, but efforts to monitor them are hampered by the irregularity and relative unpredictability of their movements. We developed a bird migration forecast system with continental scope by leveraging 23 years of spring observations to learn associations between atmospheric conditions and bird migration intensity. Our models explained up to 81% of variation in migration intensity across the United States at altitudes of 0-3000 m, and performance remained high when forecasting events 24-72 h into the future (68-72% variation explained). We infer that avian migratory movements across the United States frequently exceed 200 million individuals per night and exceed 500 million individuals per night during peak passage. Accurately forecasting bird migration will allow stakeholders to reduce collisions with illuminated buildings, airplanes, and wind turbines, predict movements under climate change scenarios, and engage the public.

---

Billions of birds migrate to distant breeding and wintering sites each year through landscapes and airspaces that are increasingly transformed by humans. Hundreds of millions die annually from anthropogenic hazards such as collisions with buildings, automobiles, and energy installations (1), and these effects are exacerbated by light pollution (2). Bird migration is characterized by dramatic pulses of movement interspersed with stopover periods of relative calm (e.g. 3, 4). Targeted efforts to reduce negative impacts on birds in flight (e.g. turning off lights and wind turbines at strategic times (5)) would be most effective if they focused on the few nights with large migration events. However, bird movements are challenging to accurately predict days or even hours in advance.

For decades, scientists have attempted to understand the drivers of avian migration. It is well known that wind, temperature, barometric pressure, and precipitation play key roles (6–8). However, it has been challenging to translate general associations into migration forecasts for continental extents while also being accurate at fine spatial and temporal resolutions (e.g. 3, 9, 10). General relationships between conditions and migration intensity are modified by local topography, regional geography, time of season, and finer properties of the atmosphere (11). Furthermore, hundreds of species with remarkably diverse behaviors frequently pass through a single location during the migration season. This complexity makes predicting continental bird migration at the assemblage level a grand challenge, requiring the integration of large environmental and behavioral datasets with methods that can capture complex associations (12).

Amassing behavioral data that appropriately characterize the spatial and temporal scale of continental bird migration has been a primary impediment. Doppler radar, which has been used across the world as a tool to study animal migration (4, 13–17), offers a realistic solution to characterize the system-wide behaviors of hundreds of migratory species (18). In the continental United States, the NEXRAD radar network comprises 143 weather surveillance radars (19), and its data archive contains two decades of continuous observation. Although designed for meteorology, these radars measure energy reflected by a diversity of aerial targets, including birds. However, only recently have advances in computational methods (e.g. 20) made it possible to use the entire radar archive for longitudinal studies of bird migration at large spatial scales.

Using the NEXRAD archive, we quantified 23 years (1995-2017) of spring bird migration from 143 radar stations across the US (Fig. 1). We developed a classifier to eliminate radar scans contaminated with precipitation. We then trained gradient boosted trees (21) to predict bird migration intensity from atmospheric conditions reported by the North American Regional Reanalysis (NARR) (22). Our model’s predictors were 19 variables, including zonal and meridional wind components, air temperature, barometric pressure, and relative humidity (Fig. S1), which we used to predict a cube-root transformed index of migration intensity (cm^2^ km^−3^). The cube root transform reduces skewness but is less extreme than a log-transformation, which would have given considerable weight to biologically unimportant differences between small values. We measured migration intensity in 100-m height bins up to an altitude of 3 km, allowing us to model the three-dimensional distribution of migrating birds over the continent.

**Fig. 1.**
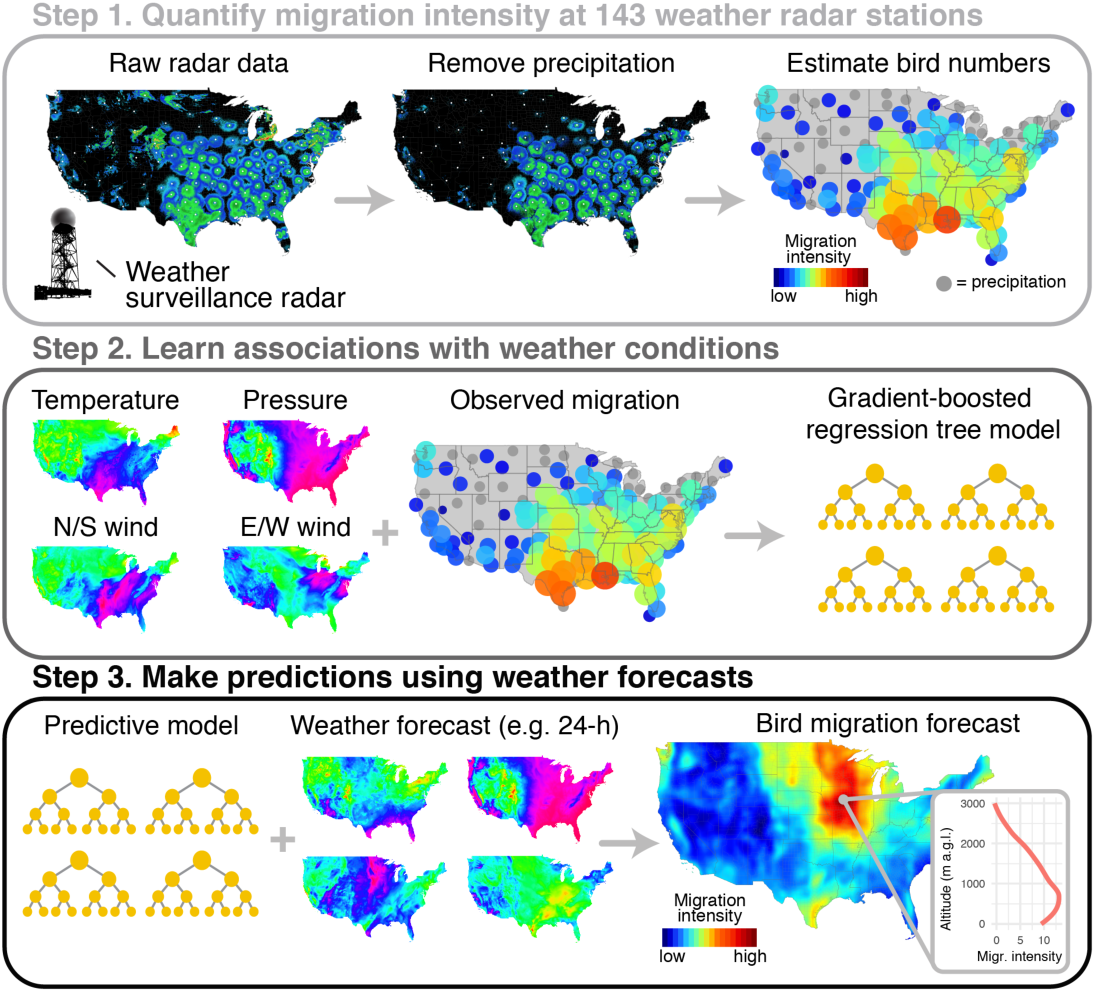
Illustration of methodology for generating migration forecasts.

Our migration forecast model explained 78.2% of variation in migration intensity over the US using instantaneous conditions information from NARR (Figs. 2 and 3B). We quantified the importance of each predictor by calculating the gain, a measure of how much a tree’s predictions improve by adding a given variable. Air temperature was the most important predictor, with an average gain nearly three times that of the next most-important predictor, date (Fig. S2). As a predictor of bird migration, temperature likely plays a dual role as an index of spring phenology (i.e. the advancing “green wave”) and a short-term signal for movement, as favorable southerly winds usually accompany warmer air masses. Migration intensity varied closely with temperature (Fig. 4); on a given day, the highest intensities occurred where temperatures were warmest (Fig. S3). This effect of temperature was also distinct from the effect of wind: given similar wind conditions, more birds took flight when temperatures were warmer (Fig. S4). Other important predictors included altitude, longitude, surface pressure, latitude, and wind (Fig. S2).

**Fig. 2.**
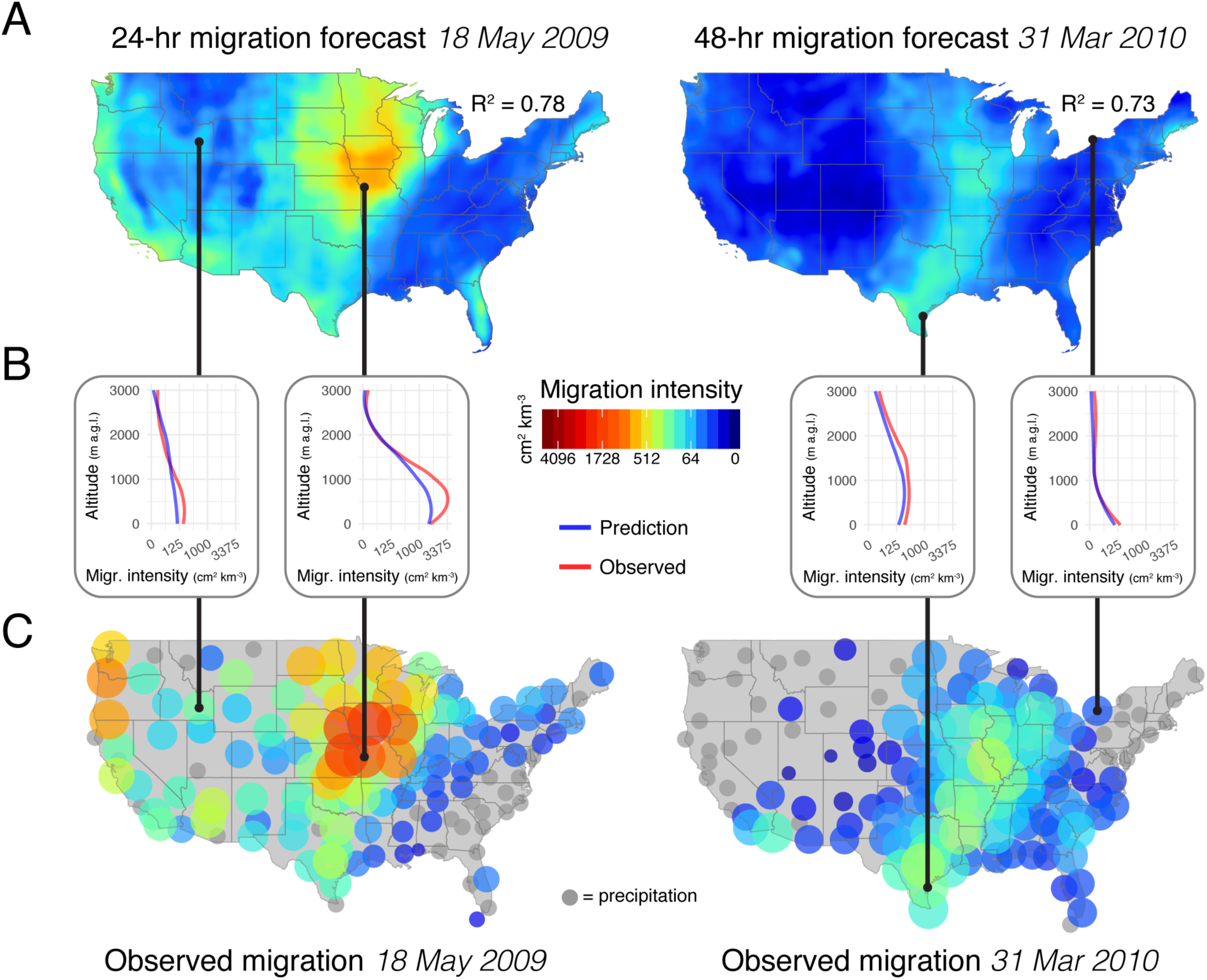
Two examples of migration forecasts (24-h and 48-h) made using test data, and their corresponding observed values. (A) Country-wide migration forecast surfaces showing the predicted mean migration intensity across altitudes. (B) Altitudinal profiles at four stations, showing predicted and observed intensity values. (C) Mean migration intensity observed at all radar stations. Gray circles show stations where migration intensity could not be measured due to the presence of precipitation.

**Fig. 3.**
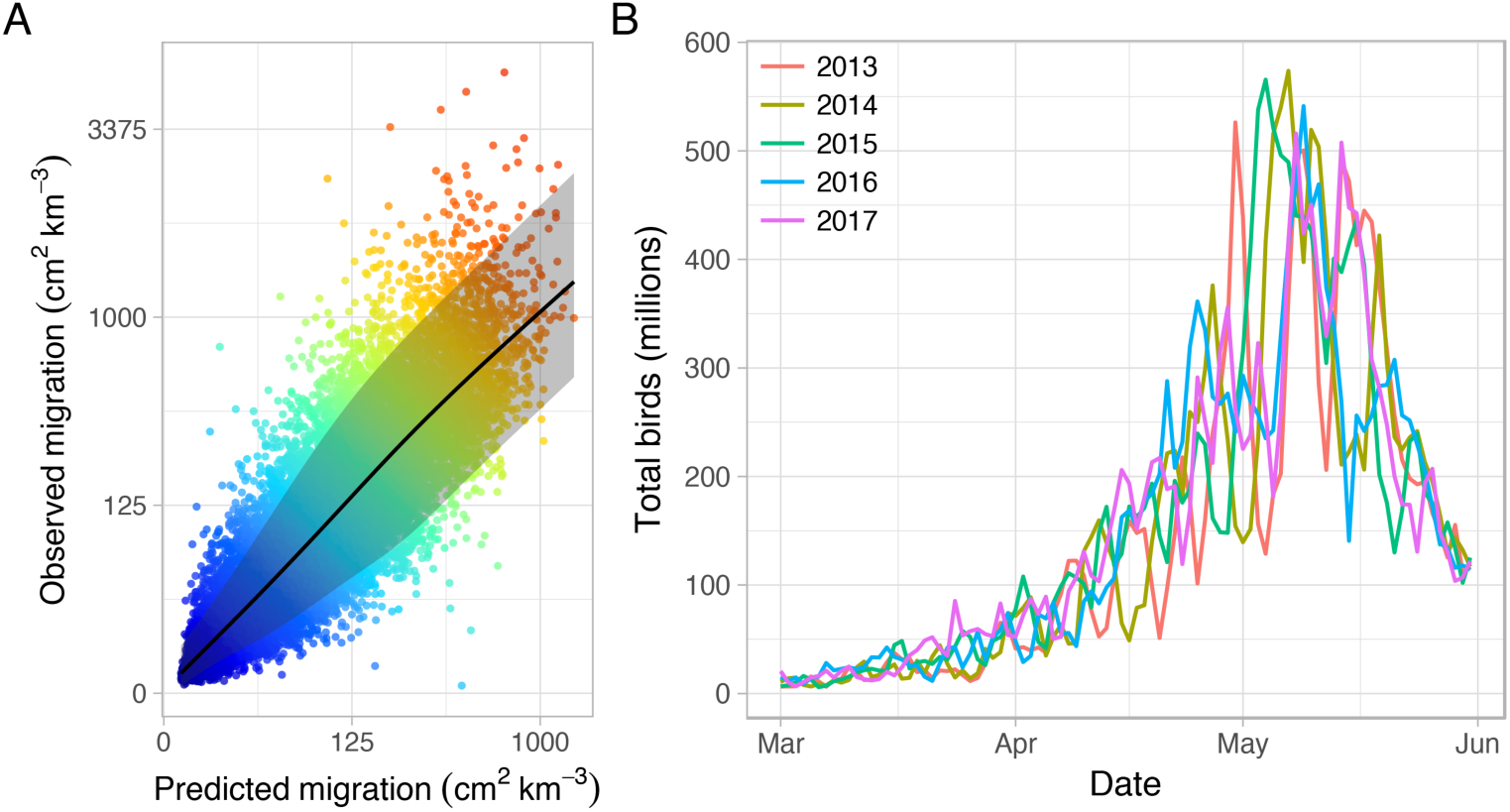
(A) Mean predicted migration intensity versus observed migration intensity for test data, with points colored by observed intensity. The scatterplot shows values after averaging across altitudes. R^2^ for the full dataset (before averaging) was 0.78. Shading shows empirical 90% prediction intervals, which covered 91.0% of observed values. (B) Nightly peak migration magnitude estimated across the continental United States for 2013-2017. Numbers of individuals estimated using a cross-sectional area per bird of 11 cm^2^ (*14*). Note that the size of migratory movements varied dramatically from night to night during the peak of the migration season.

**Fig. 4.**
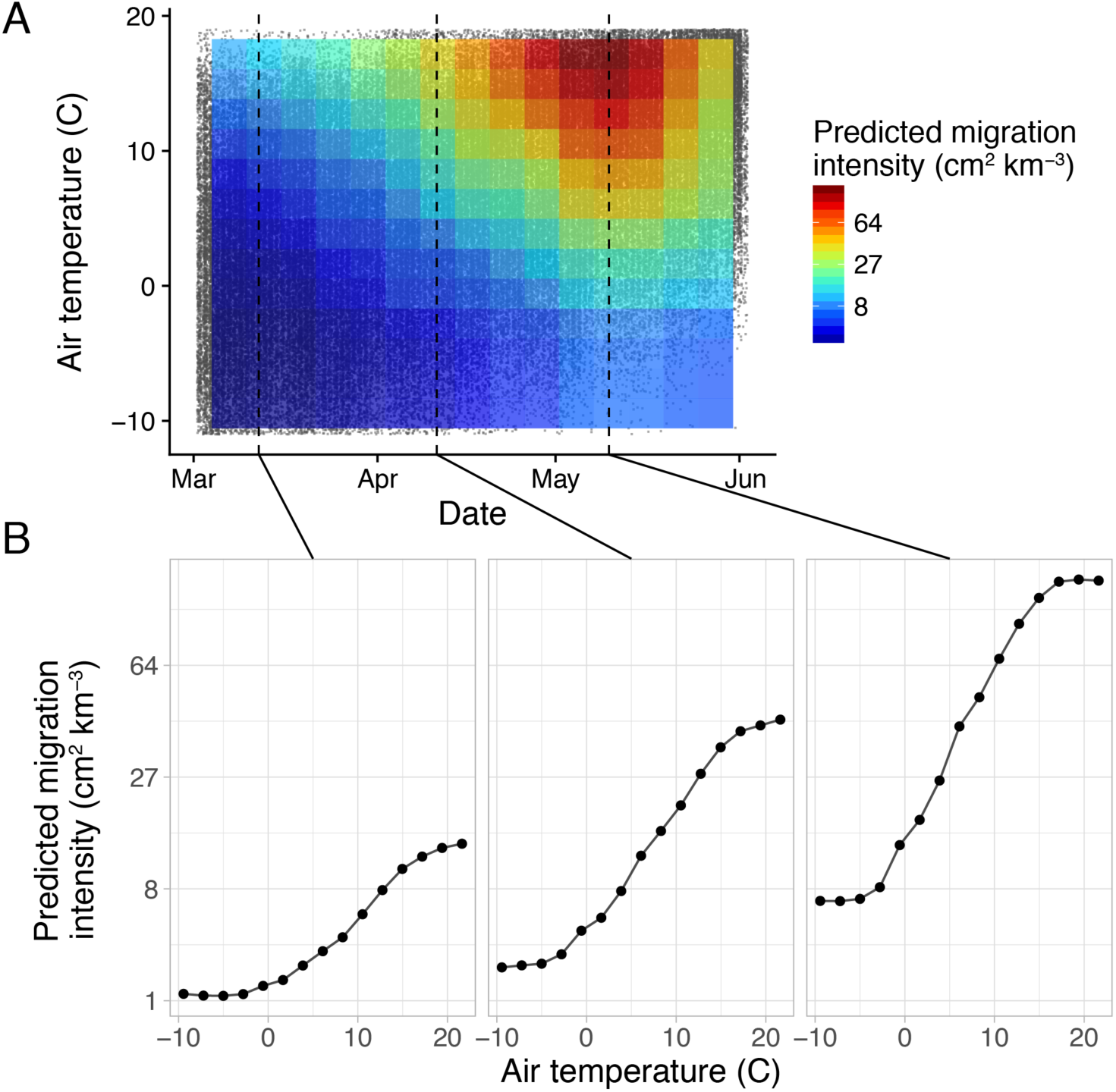
(A) Model predictions of migration intensity versus air temperature and date, the two most important predictors. Each data point on the scatterplot beneath the heatmap represents one night from one radar. Only well-supported predictions and corresponding data points shown (the outer 10% of temperature and date values are excluded). (B) For a given date, the model predicts migration intensity to vary closely with temperature.

Next, we asked how far in advance our model could accurately forecast bird migration. To achieve this, we processed archived weather forecasts from the North American Mesoscale Forecast System (NAM). Migration predictions based on NAM could form the basis of an operational migration forecast product. We evaluated our model’s performance with 24-h, 48-h, and 72-h NAM weather forecasts, expecting performance to degrade with time due to the decreasing accuracy of longer-range weather forecasts. Indeed, performance decreased for more distant forecasts, but our model still explained the vast majority of variation in observed migration intensity when using NAM for prediction (e.g. 72% variation explained with 24-h forecast) (Fig. S5). Even 72-h forecasts explained 68% of variation in the test dataset. The relatively small decline in accuracy from 24 to 72 h suggests that even longer-range forecasts may perform similarly.

To examine our ability to accurately predict migration over areas without nearby radar coverage, we iteratively removed the data from each radar station, retrained the model on the remaining data, and tested predictive performance on data from the withheld station. For this analysis, we increased the learning rate parameter of the model to make computation tractable. Compared to validation benchmarks of RMSE=1.424 (root mean square error; smaller values indicate more accurate predictions) and R^2^ = 0.756, the median RMSE and R^2^ for withheld stations were 1.426 and 0.680, respectively (Fig. S6). For 75% of withheld stations, R^2^ was 0.52 or higher. We therefore conclude that the model captures patterns of bird migration across the United States with high spatial accuracy, particularly in the Central and Eastern regions (Fig. S7); spatial variation in performance likely stems from local influences on migratory behavior (e.g. topography), which our model did not explicitly incorporate.

Previous research has suggested that migration behavior and weather conditions in the days immediately preceding a focal day can predict migration intensity (e.g. 10). We found that including atmospheric data from the preceding night and 24-h changes in conditions did improve performance, but not dramatically. Compared to our original specification (R^2^ = 0.782, RMSE = 1.318), a model that included atmospheric conditions 24 h before the focal time explained 79.5% of variation in the test data (RMSE = 1.279). Further including observed migration intensity from the previous night increased R^2^ to 80.7% (RMSE = 1.241). Despite these improvements, we favored the original specification because it could translate a single continental weather forecast into a continental migration forecast without additional inputs.

We used our model’s predictions to estimate the total number of birds actively migrating each night across the United states, assuming a cross-sectional area per bird of 11 cm^2^ (14), typical of a medium-sized songbird. Summing predictions across the country, we infer that nightly movements frequently exceed 200 million birds (Fig. 3B). Peak passage occurred in the first half of May, when the median predicted movement size was 520 million birds per night. These are the first size estimates of nightly migratory movements on a continental scale. Our estimates are directly informed by a validated model that incorporates atmospheric conditions and geography, which is an advantage over simple interpolation between radar stations. Further, these estimates are likely conservative because our model tended to underpredict the largest observed movements (Fig. 3A).

Migration forecasts will further ecological research while aiding monitoring and mortality mitigation efforts. For example, scientists can leverage our predictive framework to investigate how future changes in environmental conditions (e.g. air temperature) may influence migratory behavior. Biologists working with migratory birds and global health workers monitoring avian borne diseases (23) can use migration forecasts to anticipate bird movements. Further integration of large citizen science datasets with radar observations will provide the means to study species-specific patterns of behavior at a large scale (12). For environmental monitoring, accurate migration predictions can inform decisions to temporarily shut down lights on skyscrapers and communication towers, elevate the cut-in speed of wind turbines, halt gas flares, and take other actions to prevent avian mortality. Our model provides predictions at 100-m altitudinal resolution, which will be valuable for aircraft pilots and operators of turbines and towers. Finally, educators and birdwatchers can use migration forecasts to connect themselves and others with one of the world’s most impressive natural phenomena.

## Methods

### Doppler radar

We used the United States Next Generation Radar (NEXRAD) network to characterize spring migratory movements (March 1st to May 31st) from 1995 to 2017. These radars scan 360° at multiple elevation angles (e.g. 0.5°, 1.5°… 4.5°), fully sampling the airspace every 5 to 10 minutes. We downloaded radar scans from Amazon Web Services (https://s3.amazonaws.com/noaa-nexrad-level2/index.html), selecting those in a 30-minute window centered on three hours after local sunset. We processed scans using the WSRLIB software package (24). To characterize migration intensity, we used radar reflectivity factor, a measure of reflectance to the radar. To sample the airspace from 0-3 km above ground level, we extracted radar reflectivity factor values 5-37.5km from each radar (25) and cast them into vertical profiles with 100-m altitudinal resolution. We converted radar reflectivity factor (dBZ) to radar reflectivity (dBη) using the equation η[dB] = Z[dBZ] + β, where β = ^10log 10 (10^3^π^5^|Km|^2^/λ^4^). We set the radar wavelength (λ) to 10.71 cm, the average for NEXRAD radars and set the refractive index (|Km|^2^) to 0.93 for^ liquid water. This yielded β = 13.35. We converted dBη to η using the equation η= 10^dBη/10^, yielding units of cm^2^km^−3^. To estimate numbers of birds from η, we selected a cross section (σ) of 11cm^2^, a representative average cross-section across all bird species within a migratory season (*14*). Dividing η by σ resulted in units of birds km^−3^.

To mitigate the influence of time-invariant clutter (e.g., buildings, terrain, wind turbines), we applied binary clutter masks to each low elevation scan prior to the construction of the vertical profile of migrant intensity. Masks were generated by summing a minimum of 100 low elevation scans (0.5° elevation), starting on January 1st (16:00 UTC to 18:00 UTC) and continuing to January 15th. This time window falls well outside typical migration periods. If 100 samples were not tallied by January 15th the window of selection was expanded until the threshold was met. We classified any pixel above the 85th percentile of the summed reflectivity as clutter and masked it from our calculation of migration intensity.

To discriminate radar scans contaminated with precipitation from those containing only clear air or bird-dominated signal (hereafter termed “clear”), we created a random forest classifier using the package “randomForest” (26). We trained the classifier on 157,279 manually classified nocturnal scans selected from a 3-hour period on March 15th, April 15th, and May 15th for every radar and every year in the training set. We designed this sampling to capture any geographic, seasonal, and longitudinal patterns in the data. We extracted derived predictor variables from profiles of radar reflectivity, groundspeed, migrant track, and summaries of the number of density values above 35 dBZ (extreme densities characteristic of intense precipitation). We generated 1,000 trees and set the minimum terminal node size to 50. Overall, the model showed a 5.6% classification error. We used the algorithm to classify a total of 979,326 scans. As an additional step to reduce the inclusion of precipitation incorrectly classified as clear, we used only scans with a probability of being clear of 70% or higher (rather than a majority rule, i.e. >50%).

### Weather reanalysis

The North American Regional Reanalysis, or NARR (22), compiles data from numerous observational data sources to produce a best estimate of weather conditions that occurred in North America. The reanalysis is published in 3-hour intervals across a 32-km grid. We downloaded NARR data for 1995-2017 and extracted the following parameters at pressure levels from the surface to 600 mb: geopotential height (gpm), zonal and meridional wind components (m/s), temperature (K), relative humidity (%), vertical velocity (Pa/s), turbulent kinetic energy (J/kg), total cloud cover (%), visibility (m), albedo (%), total precipitation (kg/m2), mean sea level pressure (Pa), convective available potential energy (J/kg), and snow cover (%). To match weather data to radar stations, we averaged data within 37.5 km of each radar station. We then calculated height above ground level by subtracting surface geopotential height from the geopotential height at each pressure level, and we matched radar observations to weather data using nearest neighbor interpolation (i.e. taking the weather observation closest in time and altitude for each radar bin). Pairwise correlations among predictor variables were generally low (Fig. S1).

### Weather forecasts

The North American Mesoscale Forecast System, or NAM (https://www.ncdc.noaa.gov/data-access/model-data/model-datasets/north-american-mesoscale-forecast-system-nam), generates weather forecasts out to 84 hours; forecasts are hourly from 1-36 hours and subsequently every 3 hours until hour 84. Forecast models are run every 6 hours. NAM predictions are made on a 12-km grid. We downloaded 0Z NAM forecast data for 2008-2017, extracted the same parameters as for NARR, and matched NAM data to radar stations in the same manner as for NARR.

### Supervised learning

We used gradient boosted trees to predict bird migration from weather data (Fig. 1). We used the R implementation of XGBoost (21, 27), a highly efficient and scalable gradient boosting framework. The algorithm automatically detects nonlinear effects and complex interactions among predictors, and it is not hindered by predictor collinearity. We trained an XGBoost model on NARR weather data, with the cube root of bird density as our response variable.

We divided our dataset into three groups: a training set, for learning; a validation set, for hyperparameter tuning; and a test set, to evaluate performance. We split the dataset by whole days instead of individual data points to prevent any spatial autocorrelation from inflating performance metrics. From 2,115 total days (comprising 5,272,618 altitude bins across 143 radar stations), we randomly selected 75% of days for training, 10% for validation, and 15% for testing.

We tuned model hyperparameters with grid searches across hyperparameter space. For our first grid search, we varied maximum tree depth max_depth (10–16) and learning rate eta (0.05, 0.10, or 0.20), as these hyperparameters generally have the largest effect on performance. In all cases, we used the early_stopping_rounds argument to stop the algorithm after 10 boosting iterations in which performance on the validation set failed to improve. After selecting the values of maximum tree depth and learning rate that resulted in the best performance on the validation set, we tested the following modifications to additional parameters with a second search: decreasing subsample from 1.0 to 0.70, increasing min_child_weight from 1 to 5, and increasing gamma from 0 to 1, 10, or 30. We tried all 16 combinations of these modifications. Using the best combination of hyperparameters, we further lowered the learning rate to 0.01 and set early_stopping_rounds to 50 to determine the optimal number of boosting iterations for that learning rate. With this information, we fit a final model with learning rate = 0.01 on the combined training and validation sets. We then evaluated its performance on the test dataset (15% of data), which had been withheld from all training. To assess performance, we calculated two metrics: root mean square error and the coefficient of variation (or R^2^). We found R^2^ by first calculating the relative squared error (the sum of squared errors divided by the sum of squares explained by the mean value of the response), and then subtracting this value from 1. Therefore, an R^2^ value of 0 indicates that the model does not explain the observed data any better than a simple null model of the mean value of the observed data, while a negative R^2^ value indicates that the model explains the data worse than this null model.

We trained and tested two further modifications to the final model: one which also included additional conditions variables from the previous night (wind, temperature, surface pressure, precipitation) and their 24-h changes, and another which included these lagged weather variables plus migration intensity measured during the previous night. Our aim here was to determine how much additional explanatory power we could achieve with a model that takes into account recently observed conditions and behavior.

### Assessing performance

To assess performance of the final model using weather forecasts instead of NARR (i.e. reanalysis) data, we tested the model using archived NAM forecasts made 24 h, 48 h, and 72 h in advance.

To assess model performance at unobserved spatial locations, we performed a cross-validation where we randomly removed one station (out of 143 total) from the dataset, retrained the model on the remaining data, and tested its performance on the withheld station. To speed up computation, we increased the learning rate hyperparameter to 0.20 and maximum tree depth to 14; this resulted in a small decrease in overall performance (<2%) but sped up computation by an order of magnitude, allowing us to compare performance at all 143 sites.

### Predictor importance and partial dependencies

We identified the predictor variables that were most important for model predictions using variable importance metrics calculated by the xgboost package. Specifically, we calculated the gain, which measures how much a tree improves by adding a split on a given variable and is therefore a measure of the variable’s importance in making accurate predictions. We also generated partial dependence plots using the R package mlr (28) to explore how these variables influence predictions. As for the station cross-validation procedure, we increased the learning rate and tree depth to make computation tractable.

### Prediction intervals

We constructed empirical prediction intervals using residuals from XGBoost predictions for the validation dataset. We fitted a generalized additive model (29) on squared XGBoost residuals against the XGBoost-predicted value to account for an error variance that increased with the magnitude of the predicted value. The generalized additive model produced an estimated error variance for each predicted value, which we used to construct 90% prediction intervals using 0.05/0.95 Gaussian quantiles. We constructed separate models for upper and lower limits to allow for asymmetry in the width of the interval, and we used the Gamma distribution family in the generalized additive model to constrain the predicted variances to be non-negative.

### Forecast output and estimation of nightly migration magnitude

Using our validated migration forecast model, we made predictions across the entire 12-km NAM grid. For smooth presentation, we averaged predictions across 9×9 cell blocks. We also used our model to estimate the total number of birds migrating over the continental United States each night. For this we used the NARR dataset because it is the best retrospective estimate of occurred conditions. For each 32-km NARR grid cell covering the continental United States, we multiplied the bird density estimate by the area of the cell, and we summed totals across all grid cells for each night.

## Acknowledgments

We are grateful to Andrew Farnsworth and Dan Sheldon for invaluable discussion and guidance. We also thank Wesley Hochachka, Giles Hooker, and Jacob Calvert for helpful feedback on statistical methods. Edward W. Rose Postdoctoral Fellowship supported KGH, and Marshall Aid Commemoration Commission supported BVD. Microsoft Word template from https://github.com/finkelsteinlab/BioRxiv-Template.

## Author contributions

BVD conceived of the study, performed statistical analyses, and wrote the paper; KGH performed radar analyses, shaped the study, and contributed writing.

## Supplementary Figures

**Fig. S1.**
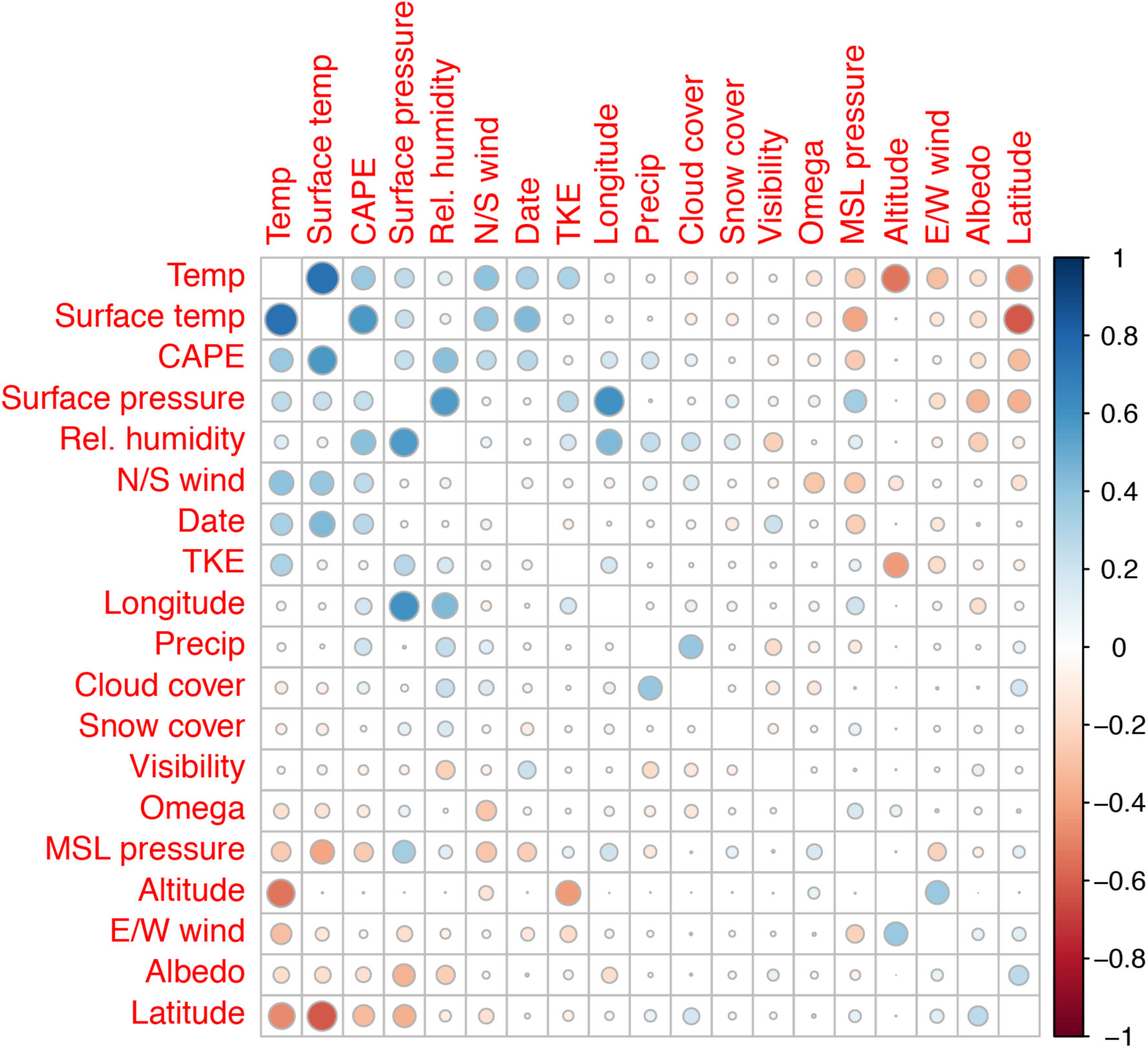
Spearman rank correlations among all pairs of predictor variables. Only one pair, Temp and Surface temp [Air temperature and surface temperature], had Spearman or Pearson correlation coefficients greater than 0.70.

**Fig. S2.**
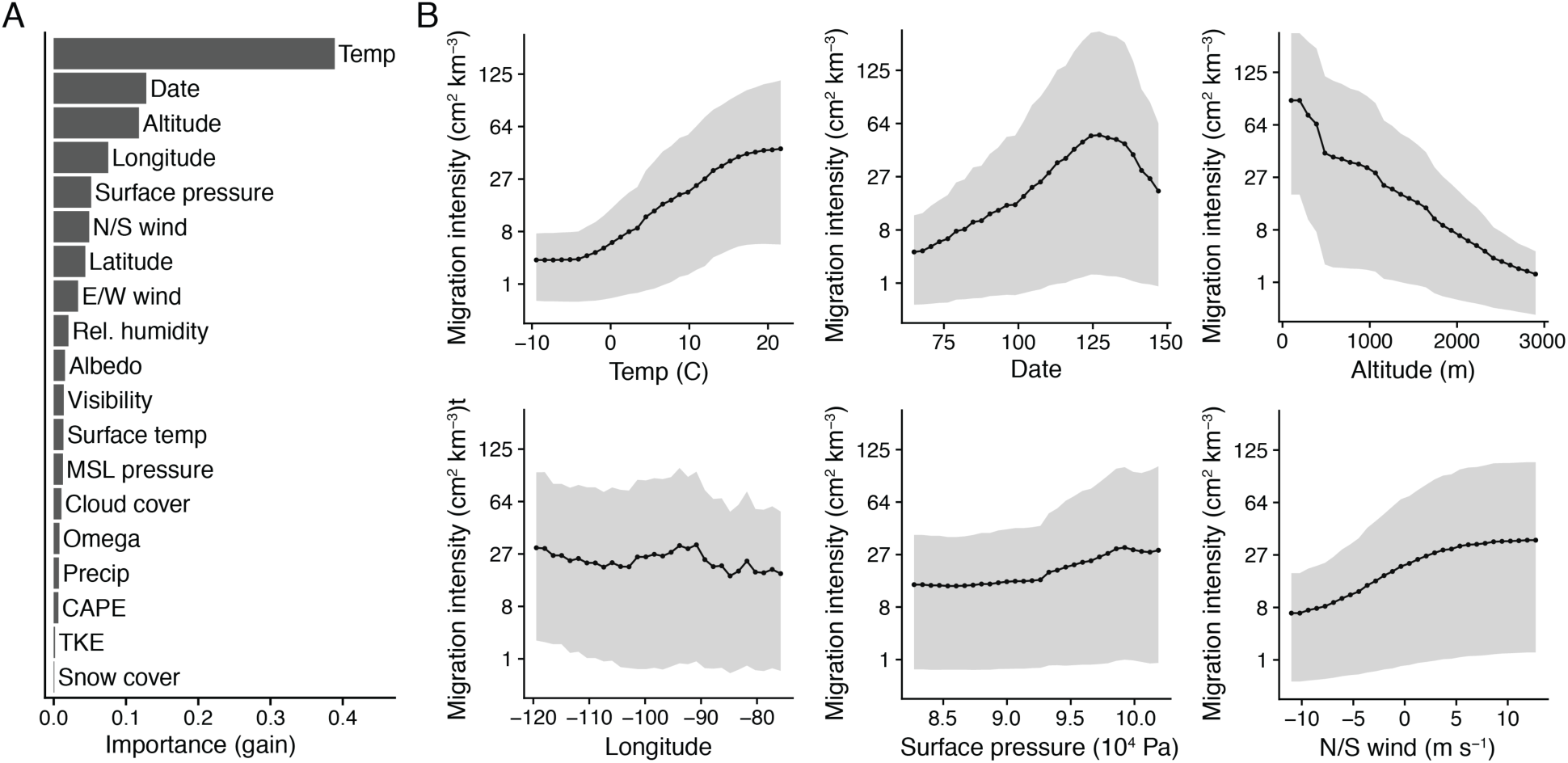
(A) Predictor importance, measured by gain. Gain is a measure of each variable’s importance in making accurate predictions. (B) One-dimensional partial dependence plots for the six most important predictor variables, evaluated for the middle 90% of predictor values. Solid lines show the mean and shading shows the middle 50% of predicted values. Narrower shading indicates that the predictor explains a greater proportion of variance in the predicted values.

**Fig. S3.**
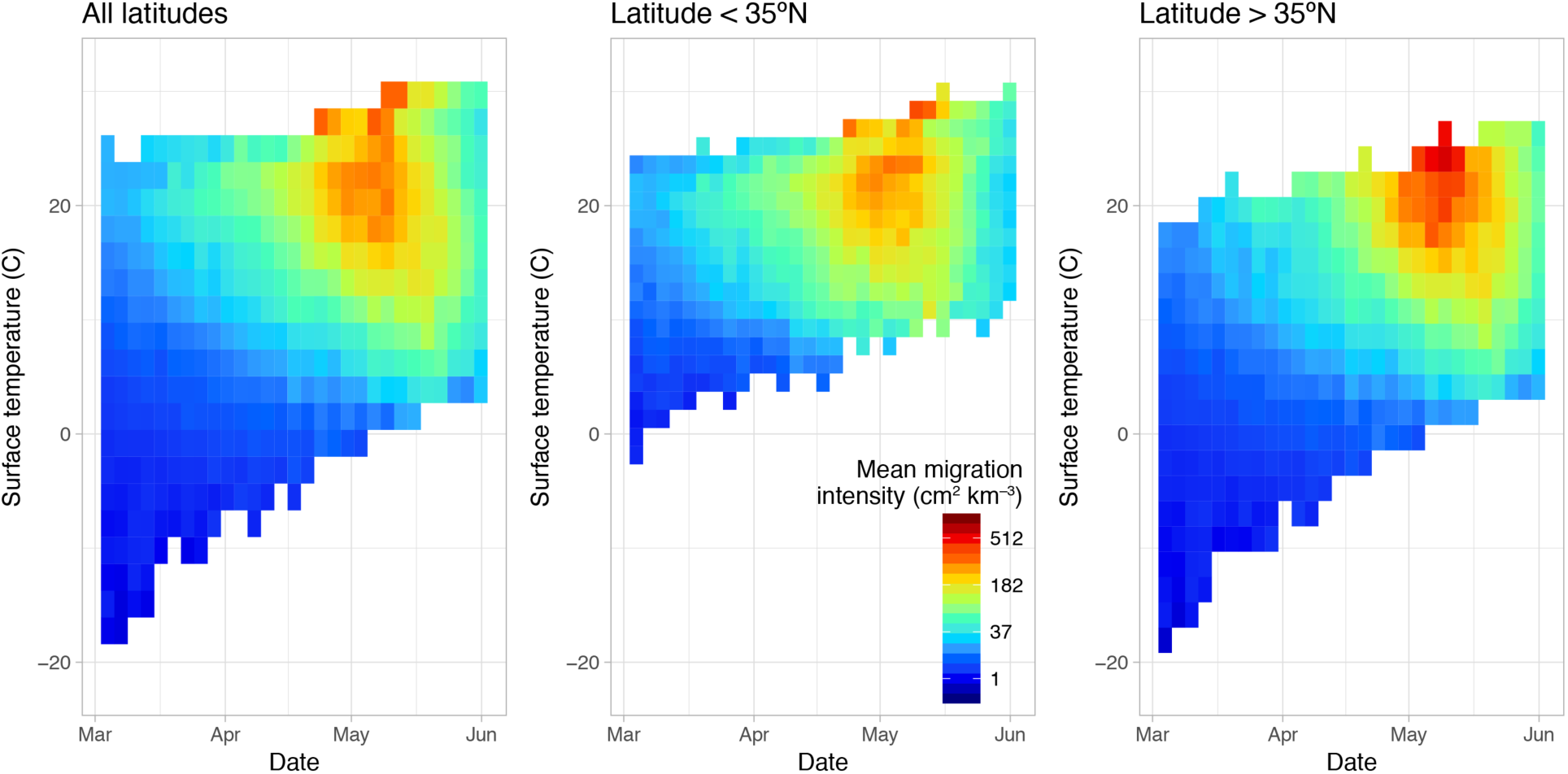
Mean observed migration intensity by date and surface temperature. For a given date, the highest migration intensities occurred where surface temperatures were warmest, especially at higher latitudes.

**Fig. S4.**
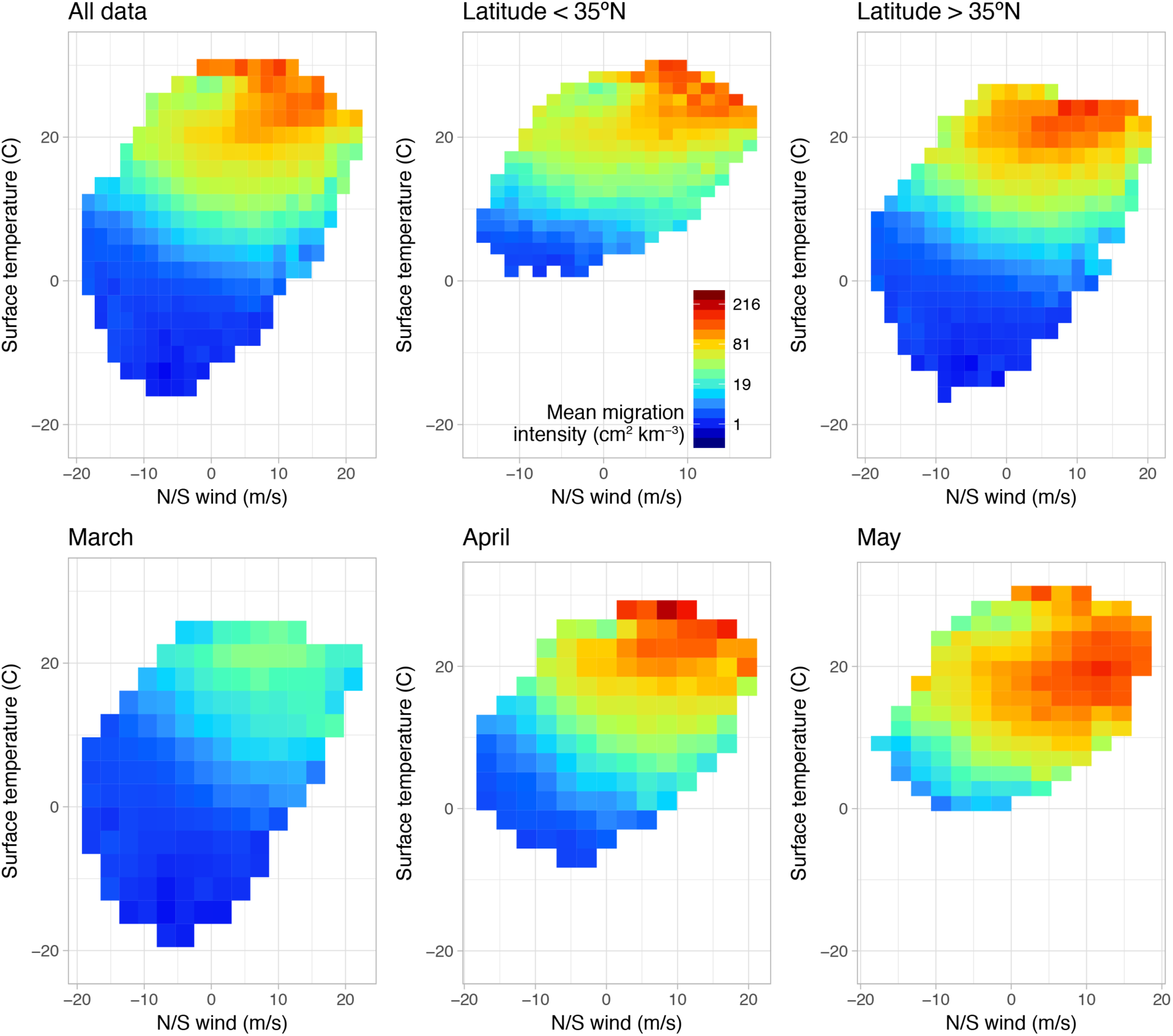
Mean observed migration intensity by surface temperature and wind direction.

**Fig. S5.**
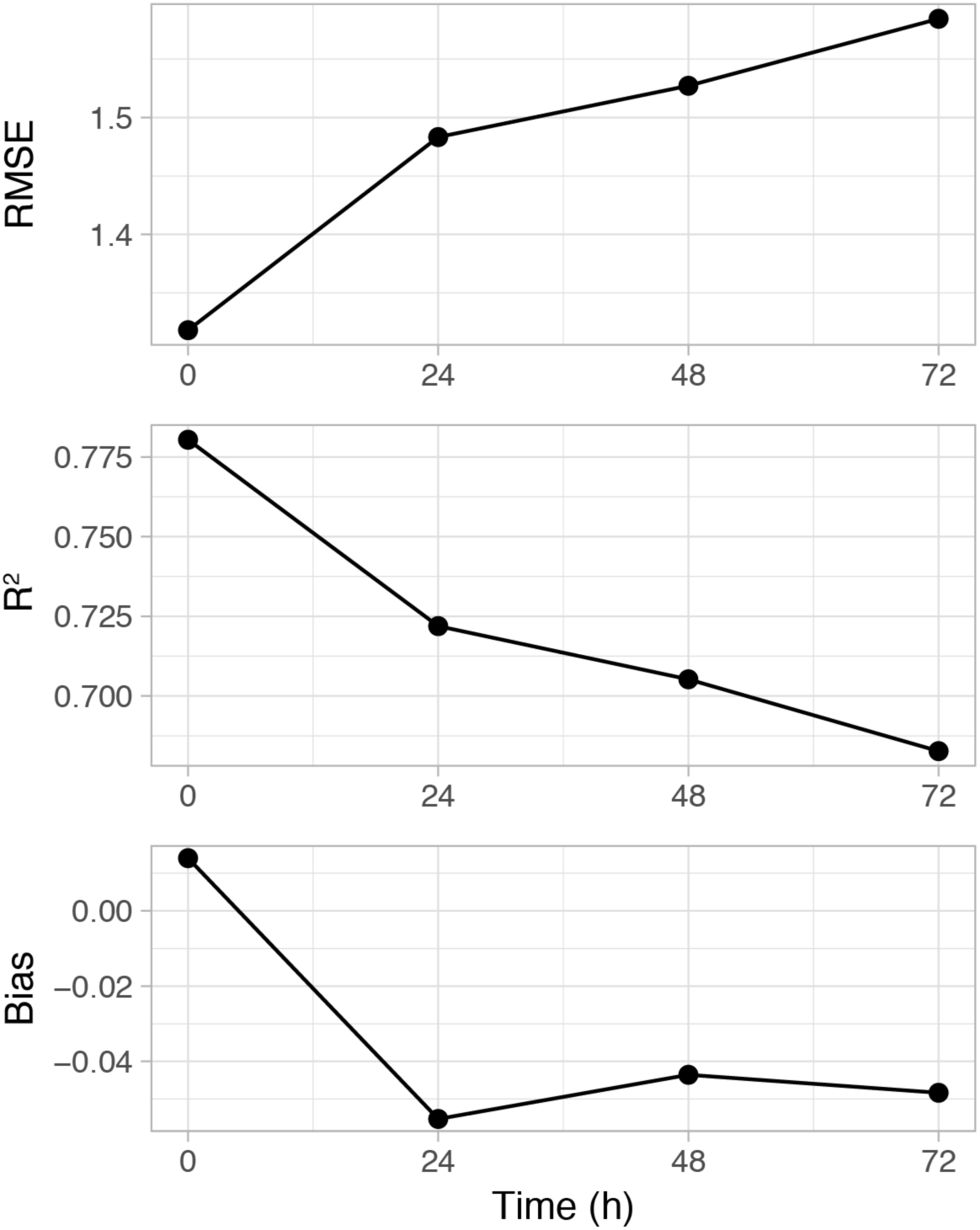
Model performance using NAM weather forecast data compared to NARR reanalysis data.

**Fig. S6.**
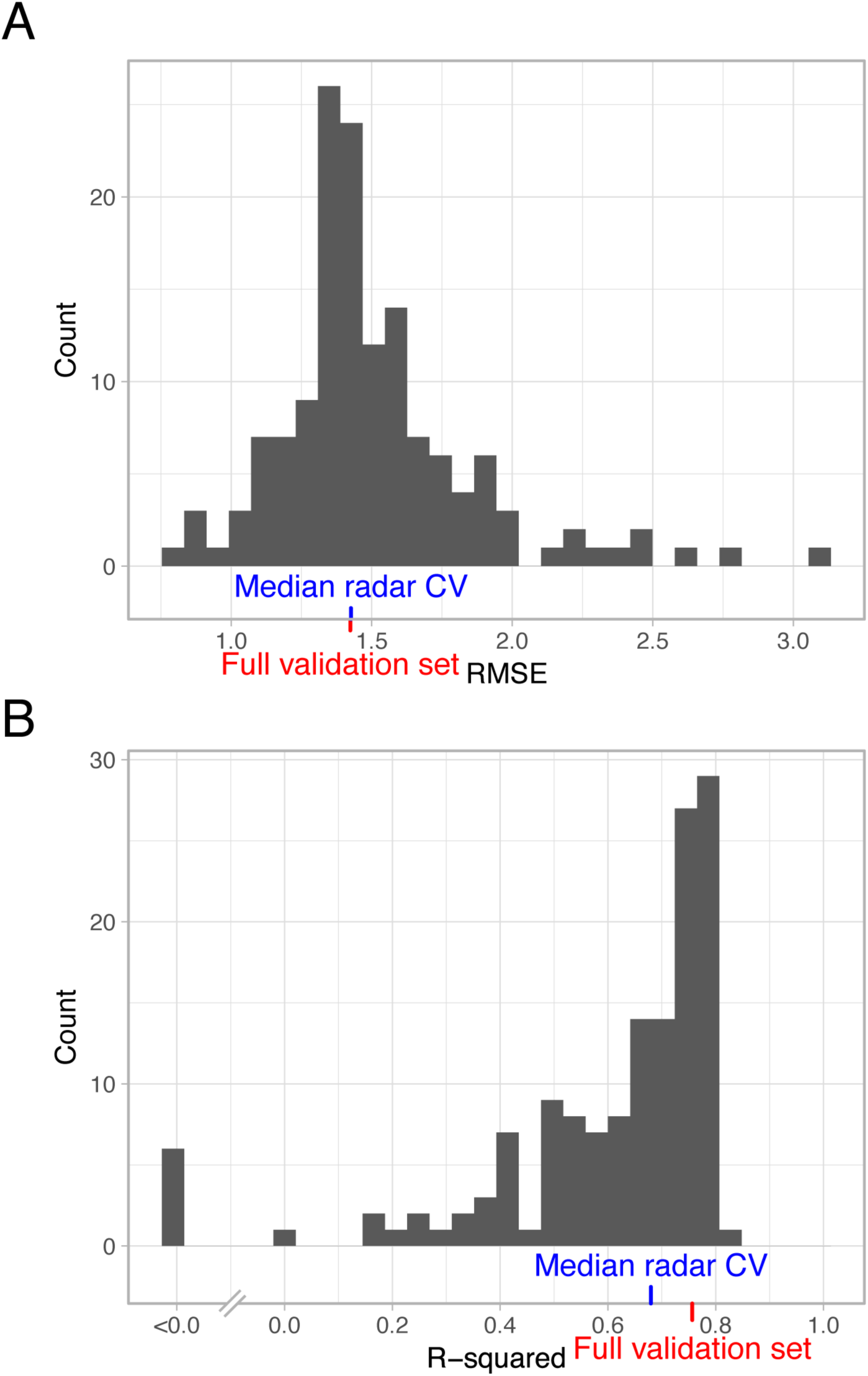
Model performance at unobserved locations. Histograms show the distribution of performance metrics for radar stations that were withheld from the training dataset. The blue tick marks show the median value across sites, and the red tick marks show the corresponding value for the randomly-selected validation set (all locations included).

**Fig. S7.**
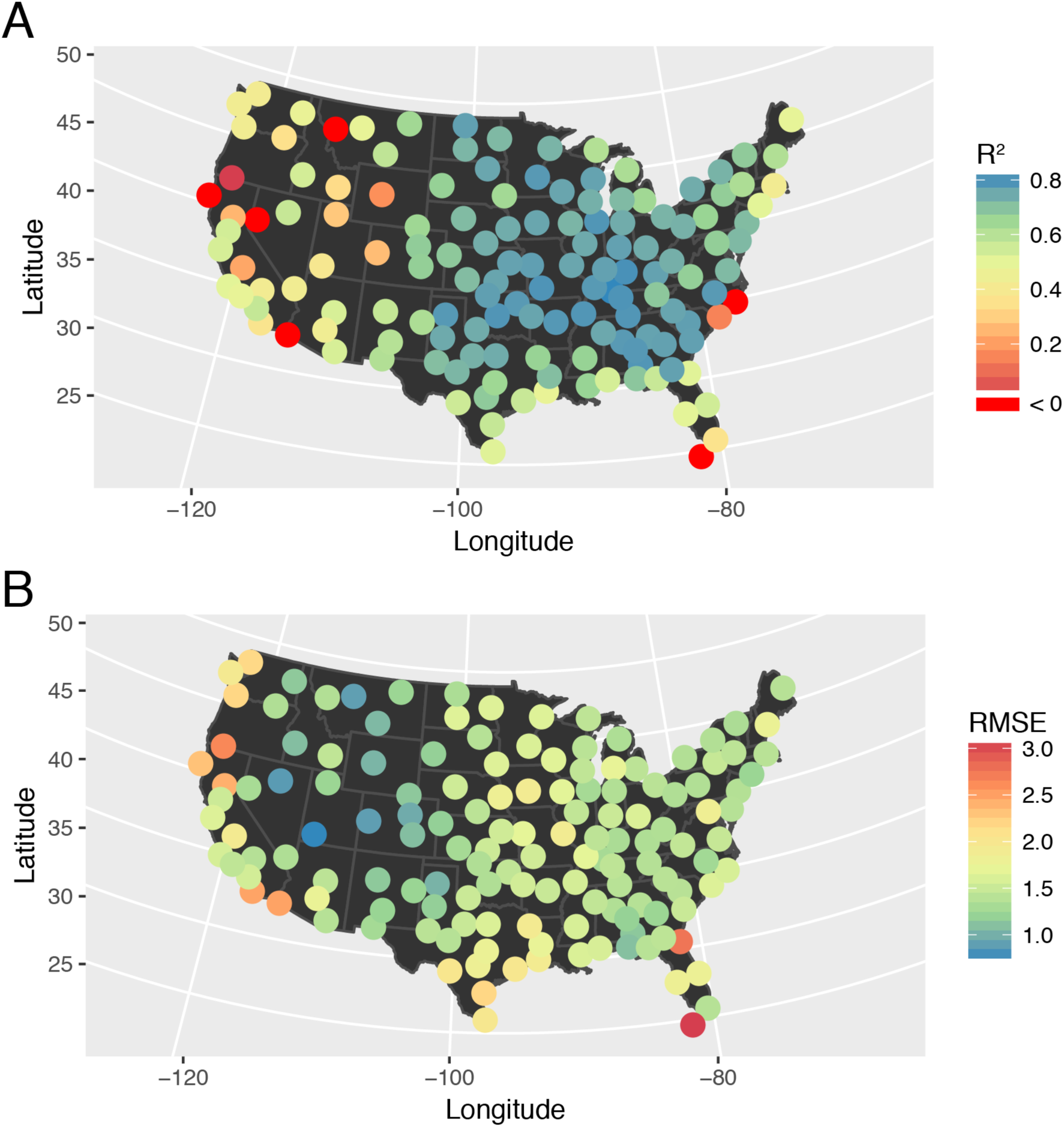
(A) Relative and (B) absolute performance at radar stations withheld from the training dataset. Performance was best at interior sites, especially in the Central and Eastern United States. Relative performance was poor (R^2^ < 0) at small minority (4%) of withheld sites, which may be due to local influences such as topography (e.g. see Florida Keys). However, at most of these sites, absolute performance (RMSE) was not substantially worse than at nearby locations.

